# Fine-tuning of a CRISPRi screen in the seventh pandemic *Vibrio cholerae*

**DOI:** 10.1101/2024.07.03.601881

**Authors:** Kevin Debatisse, Théophile Niault, Sarah Peeters, Amandine Maire, Baptiste Darracq, Zeynep Baharoglu, David Bikard, Didier Mazel, Céline Loot

## Abstract

*Vibrio cholerae O1 El Tor*, the etiological agent responsible for the last cholera pandemic, has become a well-established model organism for which some genetic tools exist. While CRISPRi has been applied in *V. cholerae*, improvements were necessary to upscale it and enable pooled screening by high-throughput sequencing in this bacterium. In this study, we introduce a pooled genome wide CRISPRi library construction specifically optimized for this *V. cholerae* strain, characterized by minimal cytotoxicity and streamlined experimental setup. This library allows the depletion of 3, 674 (98.9%) annotated genes from the *V. cholerae* genome. To confirm its effectiveness, we screened for essential genes during exponential growth in rich medium and identified 368 genes for which guides were significantly depleted from the library (log2FC < - 2). Remarkably, 82% of these genes had previously been described as hypothetical essential genes in *V. cholerae* or in a closely related bacterium, *V. natriegens*. We thus validated the robustness and accuracy of our CRISPRi-based approach for assessing gene fitness in a given condition. Our findings highlight the efficacy of the developed CRISPRi platform as a powerful tool for high-throughput functional genomics studies of *V. cholerae*.

**GRAPHICAL ABSTRACT:** 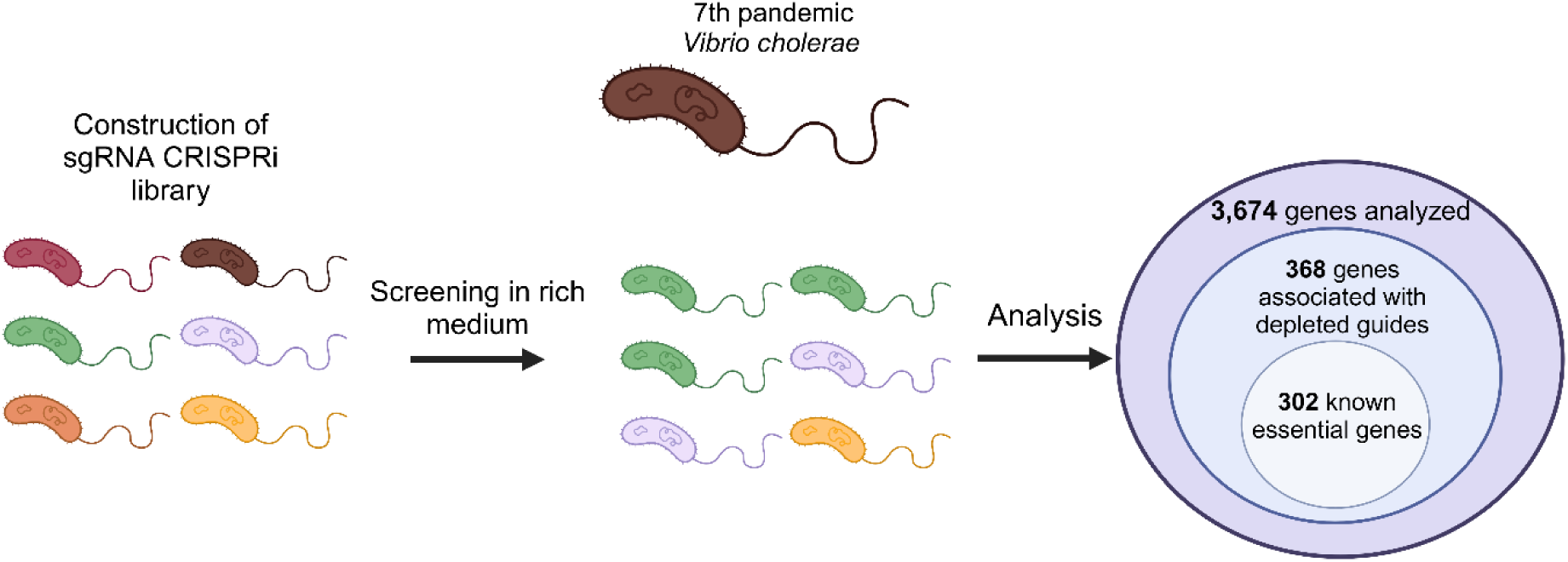

## INTRODUCTION

*Vibrio cholerae O1 El Tor* strains are the causative agents of the 7^th^ cholera pandemic (7PET), which began in 1961 and is still ongoing today (1). These strains have been extensively studied over the last few decades, and many genetic tools have been developed to better understand their biology. One of the most powerful genetic approaches is the use of global screening after construction of pooled knockout mutant libraries. The most common development called Tn-seq corresponds to random transposon insertion libraries followed by high-throughput sequencing. This approach has been used in several studies to map the list of essential genes (2, 3). However, Tn-seq presents some important limitations: (1) Complexity of library construction: following DNA extraction numerous manipulation steps are needed, which can be time-consuming and prone to experimental error. (2) Sequencing depth requirement: to obtain interpretable results, sequencing must be performed at a high depth to efficiently cover the entire bacterial genome. This also limits the multiplexing of the samples. (3) Transposon insertion bias: Transposons tend to preferentially insert into certain regions of the genome, which can introduce bias into screening results (4).

Here we report a low-cost, labor-saving approach for establishing global knockdown libraries through CRISPR interference (CRISPRi). CRISPRi is based on the use of a catalytically inactivated version of the RNA-guided nuclease Cas9 from *Streptococcus pyogenes* called dCas9. While dCas9 lacks the capacity to cleave DNA strands, it retains its ability to recognize its target DNA when paired with its associated single-guide RNA (sgRNA) thus blocking the progression of RNA synthesis (5, 6). As for Cas9 activity, the only requirement for dCas9 binding is the presence of a short DNA motif termed a <u>p</u>rotospacer <u>a</u>djacent <u>m</u>otif (PAM), typically 5’-NGG, located next to the 20-nucleotide target sequence.

CRISPRi screens have provided valuable insights in microbiology by the development of two distinct strategies, arrayed (ordered clones) or pooled screens (7). Arrayed CRISPRi enables the assessment of individual knockdown phenotypes but is limited by scaling constraints. Conversely, pooled screens operate on a larger scale and have been extensively and successfully used in a large set of bacteria (*E. coli* (8–12), *Bacillus subtilis* (13), *Mycobacterium tuberculosis* (14)*, Staphylococcus aureus* (15), *Streptococcus pneumoniae* (16), *Synechocystis* (17) and *Vibrio natriegens* (18), to cite a few).

In this study, we developed a platform for dCas9 and sgRNA expression optimized for the representative 7PET strain N16961. This platform corresponds to a single mobilizable plasmid encoding two CRISPRi functional units. In our setup, the sgRNA expression is driven by a constitutive promoter while a *dCas9* thermosensitive allele named tsRC9 is placed under the control of the P_BAD_ arabinose-inducible promoter (19). Together, these two features ensure a tight control of *dCas9* expression, enabling dCas9 toxicity to be greatly reduced and gene depletion to be carried out only at the desired time. Using this platform, a library of mobilizable plasmids containing 11, 125 different guides has been constructed to potentially target and repress nearly all 3, 715 genes from this *V. cholerae* strain. This library has been transformed into a donor *E. coli* strain and can now be easily transferred to *V. cholerae* by conjugation with high efficiency. As a proof of concept, we performed a genome-wide pooled dCas9 knockdown screening in a *V. cholerae* strain during growth in rich medium (LB) for 15 generations. We then compared the results of this screen with previously published Tn-seq experiments and demonstrated a significant overlap in the list of essential genes during exponential growth. This study validates the CRISPR-dCas9 screening approach as useful and practical tool for carrying out numerous genomics studies on *Vibrio cholerae* strains.

## MATERIAL AND METHODS

### Bacterial strains, plasmids and primers

The different bacteria strains, plasmids and primers that were used in this study are described in Table 1, 2, 3 and 4.

**Table 1:** Strains used in this study

| Strain number | Relevant genotypes or description | References |
| --- | --- | --- |
| **Bacterial strains** |  |  |
| 4196 | β2163, (F-) RP4-2-Tc::Mu ΔdapA::(erm-pir) [Km<sup>R</sup> Em<sup>R</sup>] | (20) |
| 8195 | DH5α (F-) supE44 ΔlacU169 (ϕ80lacZΔM15) ΔargF hsdR17 recA1 endA1 gyrA96 thi-1 relA1 | Laboratory collection |
| 8637 | N16961 hapR+ [Str<sup>R</sup> Gm<sup>R</sup>] | Laboratory collection |
| 8725 | π3813, lacIq thi-1 supE44 endA1 recA1 hsdR17 gyrA462 zei-298::Tn10ΔthyA::(erm-pir116) [Em<sup>R</sup>] | (21) |
| 8726 | β3914, β2163 gyrA462 zei-298::Tn10 [Km<sup>R</sup> Em<sup>R</sup>] | (21) |
| K329 | 8637 Δvc2338 (lacZ) | Laboratory collection |
| S430 | A1552 ΔddmABC | (22) |
| S907 | DB3.1 ccdB<sup>R</sup> | D. Bikard lab |
| T045 | CHE02 AcrIIA4 | D. Bikard lab<br>Strain used for the construction of the gRNA library |
| T650 | S907 AcrIIA4 | This study<br>An anti-CRISPR AcrIIA4 expression cassette was cloned onto the pOSIP backbone (23) |
|  |  | (pCH3) and integrated into the HK022 *attP* site.<br>Strain used to build the vector containing *ccdB* |
| U267-U268-U278 | 8637Δ*ddmABC*::FRT-*km*-FRT | PCR from S430 with o7100/o7101 and recombined by natural transformation in 8637. |
| U322-323-324 | 8637Δ*ddmABC* | *Km* excision from U267, U268, U278 using the pMP326 flippase expressing plasmid. |
| V070-V071-V072 | 8637Δ*lacZ*Δ*ddmABC*::FRT-*km*-FRT | PCR from S430 with o7100/o7101 and recombined by natural transformation in 8637Δ*lacZ* |
| **Conjugated and transformed strains** |  |  |
| P016 | β3914 + pMP326 | This study |
| S890 | β3914 + pS890 | This study |
| S891 | β3914 + pS891 | This study |
| T025 | β3914 + pFR56 | This study |
| T026 | β3914 + pN635 | This study |
| T662 | T650 + pT312 | This study |
| U722 | T045 + gRNA library | This study |
| T922 | β2163 + pS890 | This study |
| T933 | β2163 + pS891 | This study |
| gRNA donor | β2163 + gRNA library | This study |
| gRNA Vch | U322 + gRNA library | This study |
| V036-V037-V038 | U322-U323-U324 + pS891 | This study |
| V039-V040-V041 | U322-U323-U324 + pS890 | This study |
| V051 | T045 + pV051 | This study |
| V110-111-112 | V070-V071-V072 + pS891 | This study |
| V122-V123-V124 | U322 + pV051 | This study |
| V724-V725-V726 | U322 + pV724 | This study |

**Table 2:** Plasmids used in this study

| Plasmid number | Plasmid description | Relevant properties and construction |
| --- | --- | --- |
| pBAD43 | pSC101, *araC*, P<sub>BAD</sub> promoter | *ori*<sub>pSC101</sub>, [Spec<sup>R</sup>], (24) |
| pCH3 | pOSIP AcrIIA4 | Gift from David Bikard |
| pFR56 | pFR56-random-sgRNA | *ori*<sub>p15a</sub>, [Cm<sup>R</sup>], (11) |
| pFR097 | *araC*, P<sub>BAD</sub> promoter, *tsRC9* | *ori*<sub>Cl<sub>o</sub>DF13</sub>, [Spec<sup>R</sup>], (19) |
| pMP108 | pSC101-flippase | *ori*<sub>pSC101</sub>, [Carb<sup>R</sup>], (25) |
| pMP326 | pSC101-flippase, *tsRepA* | Ts point mutation was added on pMP108 using oMV1341 and oMV1342. |
| pN635 | pFR56-*rpsL*-sgRNA | *ori*<sub>p15a</sub>, [Cm<sup>R</sup>],<br>Goldengate using pFR56 to add the *rpsL* sgRNA with annealing of |
|  |  | o6336-6337. |
| pTN127 | pFR56::tsRC9 | *ori*<sub>p15a</sub>, [Cm<sup>R</sup>], *tsRC9* gene fragment was amplified from pFR097 using oTN166/oTN167 primers. The pFR56 plasmid was amplified using oTN168/oTN169 primers. The two fragments were assembled using Gibson assembly and transformed in a DH5α. |
| pTN253 | pTN127::araC-P<sub>BAD</sub> | *ori*<sub>p15a</sub>, [Cm<sup>R</sup>], *araC*-P<sub>BAD</sub> module was amplified from pBAD43, utilizing oTN223/oTN224 primers. The pTN127 plasmid was amplified using oTN225/oTN226 primers. The two fragments were assembled using Gibson assembly and transformed in a DH5α. |
| pT312 | pTN253-*ccdB* | *ori*<sub>p15a</sub>, [Cm<sup>R</sup>], the fragment *ccdB* was amplified by PCR from pS854 using o6703/o6704 primers. The pTN253 vector was amplified by PCR using o6701/o6702 primers. Assembly of these two fragments was achieved by performing Gibson Assembly and transformed in a π3813 (8725). Vector used to construct the sgRNA library. |
| pS854 | pFR56-*ccdB* | *ori*<sub>p15a</sub>, [Cm<sup>R</sup>], *ccdB*<sup>+</sup>, (11) |
| pS890 | pTN253-*rpsL*-gRNA | *ori*<sub>p15a</sub>, [Cm<sup>R</sup>] |
| pS891 | pTN253-random-gRNA | *ori*<sub>p15a</sub>, [Cm<sup>R</sup>] |
| pV051 | pS891ΔtsRC9 | *ori*<sub>p15a</sub>, [Cm<sup>R</sup>]<br>Blunt-end ligation of a fragment obtained through PCR from pS891 with o7203/o7204 primers. |
| pV724 | pTN253-*lacZ*-gRNA | *ori*<sub>p15a</sub>, [Cm<sup>R</sup>],<br>Goldengate using pTN253 to add the *lacZ* sgRNA with annealing of o7239/o7240. |

**Table 3:**
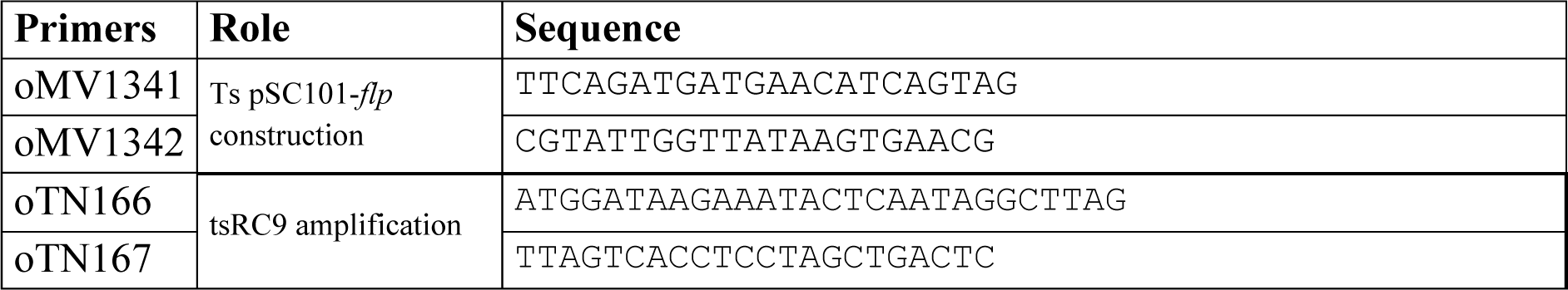
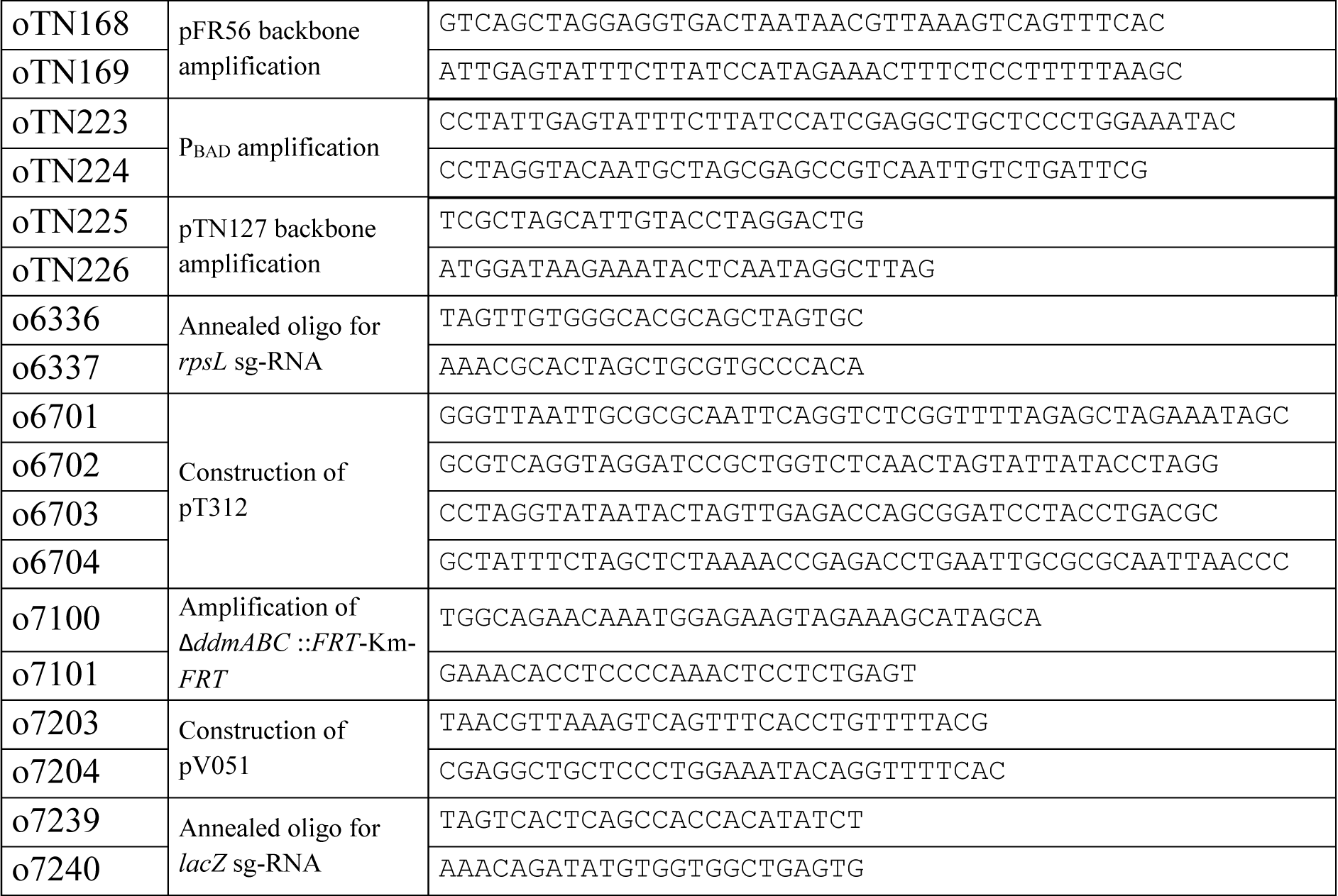
Primers used in this study

**Table 4:**
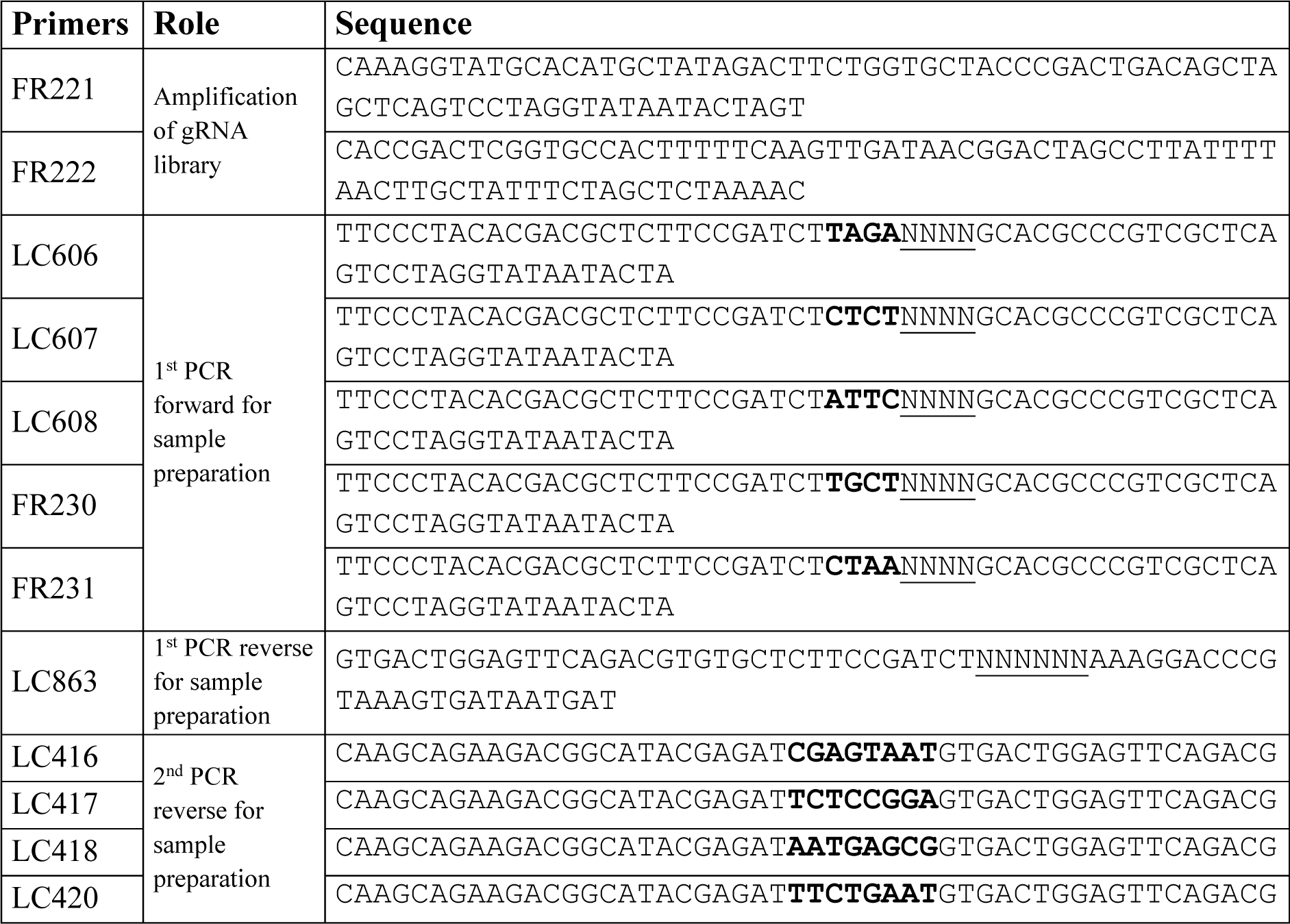
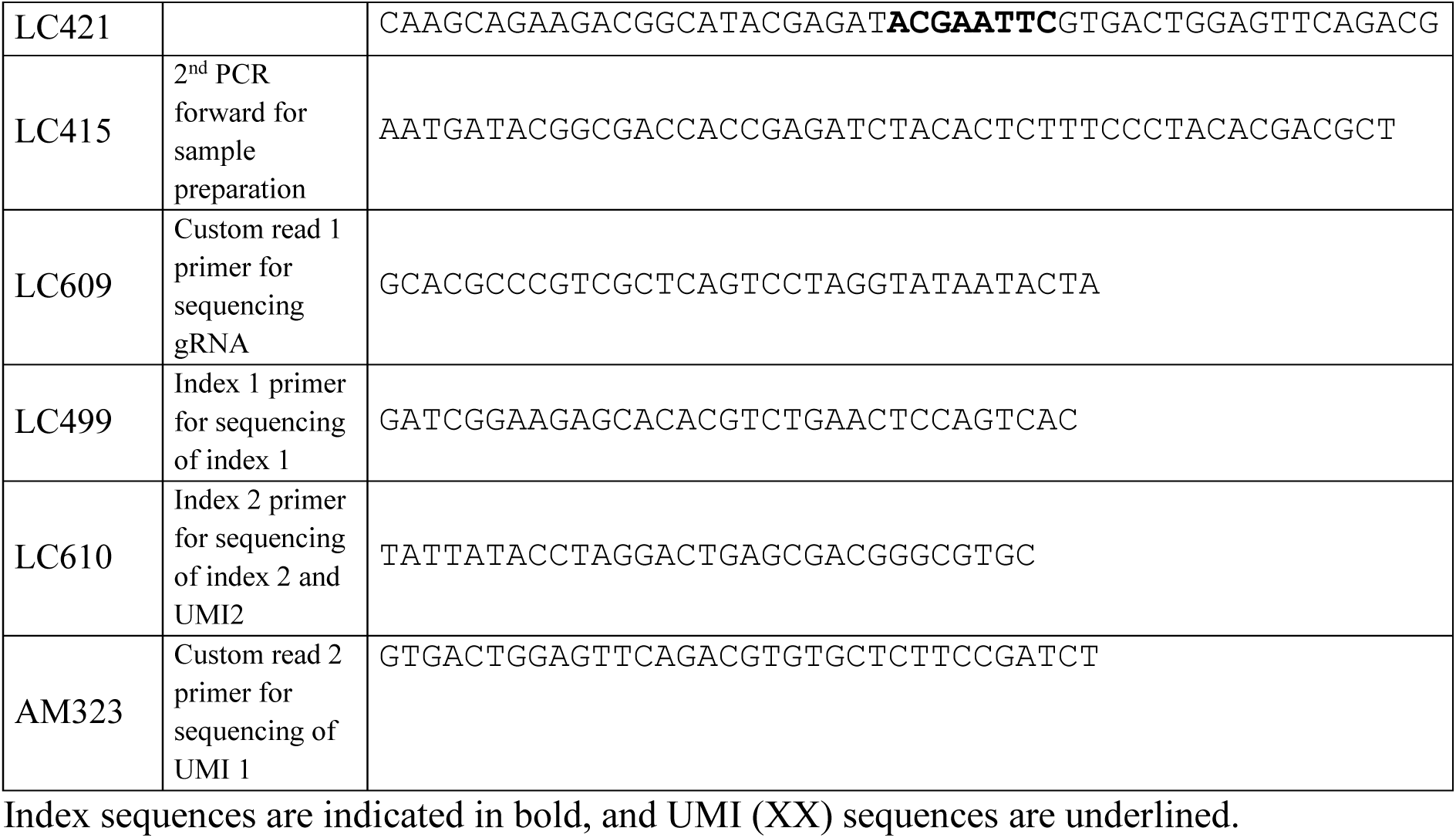
Primers used for library construction and sequencing

### Media

*Escherichia coli* and *Vibrio cholerae* were grown in Luria Bertani broth (LB) at 30°C, 37°C or 42°C. In the case of *E. coli*, antibiotics were used at the following concentrations: chloramphenicol (Cm), 25 µg/mL, kanamycin (Km), 25 µg/mL. Diaminopimelic acid (DAP) was supplemented when necessary to a final concentration of 0.3 mM while Isopropyl β-D-1-thiogalactopyranoside (IPTG) was added to reach a final concentration of 200 µg/mL. To induce the P_BAD_ promoter, L-arabinose (Ara) was added to a final concentration of 0.2 mg/mL or 2 mg/mL; to repress it, glucose (Glc) was added to a final concentration of 10 mg/mL. *V. cholerae* strains were cultivated in the same conditions and with the same antibiotic concentrations except for Cm, that was supplemented at a final concentration of 5 μg/mL.

### Strain and plasmid constructions

All PCR reactions were performed using the Phusion High-Fidelity PCR Master Mix (Thermo Scientific, Lafayette, CO, USA) or KAPA HiFi DNA polymerase (Roche), and all diagnostic PCR reactions were performed using DreamTaq DNA Polymerase (Thermo Scientific, Lafayette, CO, USA). Oligonucleotides were synthesized by Eurofins Genomics. Gibson Assembly was performed using the master mix (New England Biolabs, NEB) according to the manufacturer’s instructions. Golden Gate assembly was used to clone sgRNAs into the desired backbone using BsaI (BsaI-HF v2, New England Biolabs, NEB) (26). Plasmid DNA was extracted either using GeneJET^TM^ Plasmid Miniprep kit, or GenElute^TM^ HP Plasmid Midiprep kit (Sigma-Aldrich) or PureLink^TM^ HiPure Plasmid filter Maxiprep Kit (Thermo Scientific, Lafayette, CO, USA). Natural transformation was performed according to the procedure described in (27).

### Determining optimal induction conditions (Drop test)

*V. cholerae* mutants lacking the *ddmABC* operon (Δ*ddmABC*) and harboring the pdCas9 plasmid with either a sgRNA targeting an essential gene or a non-targeting control guide were revived from -80°C stocks by streaking onto LB agar plates supplemented with Cm and Glc and were incubated overnight at 42°C (preventing any tsRC9 leakage). Individual colonies were then inoculated in LB broth containing Cm and Glc and OD_600_ was adjusted to 0.1. The cultures were incubated at 42°C with shaking until OD_600_ reached 0.6. Cells were pelleted and resuspended in fresh LB medium to remove residual glucose. For the induction assay, serial dilutions of the bacterial culture, were spotted onto LB agar plates containing Cm to maintain plasmid selection. Three different conditions were tested for each dilution series: plates supplemented with 1% Glc to repress dCas9 expression, plates with either 0.02% or 0.2% Ara to induce dCas9 expression, and plates without any additional sugar to assess the basal expression level from the P_BAD_ promoter. The experiment was conducted in biological triplicate for each inducer concentration.

### Competition assay

To investigate competitive growth dynamics, we utilized two genetically distinct strains of *V. cholerae*. The first strain, a *lacZ*-mutant (Δ*vc2338*), harbors the pTN253 plasmid with a non-targeting sgRNA (random), this strain forms white colonies when grown on LB agar supplemented with Xgal. The second, a *lacZ*+ strain, contains pTN253 with a sgRNA targeting *rpsL* gene, this strain forms blue colonies on LB Xgal plates. Both strains were cultured independently in LB broth supplemented with Cm and Glc at 42°C until an OD_600_ of 0.6 was reached. Subsequently, equal volumes of these cultures were mixed to achieve a 1:1 ratio. The mixed culture was washed once with LB medium to eliminate residual glucose and then diluted to an OD_600_ of 0.003 in LB medium containing Cm with the following conditions: no added sugar, 1% Glc, 0.02% Ara, or 0.2% Ara. The diluted culture was incubated at 30°C for 10 bacterial generations. To quantify the competitive outcome, serial dilutions of the co-culture were diluted and spotted onto LB agar plates supplemented with Glc and X-gal, followed by overnight incubation at 42°C. The number of white and blue colonies were counted for each condition. The entire competition assay was performed in biological triplicate.

### β-galactosidase assay

To assess the level of repression that can be achieved with tsRC9 under P_BAD_, β-galactosidase activity from different cell extracts were measured. The repression level after 15 generations of growth was evaluated in strains harboring pTN253 with a guide targeting the *lacZ* reporter gene located on chromosome 1 (*vc2338*) at varying levels of induction (0.02% or 0.2%) of Ara. For the lysis, 25 µl of chloroform and 25 µl of 0.1% SDS were added to 1 mL of bacterial culture, then vigorously vortexed for 45 seconds to ensure complete mixing and left at room temperature for 5 minutes to allow the lysis reaction to take place. Next, 250 µL of the lysate was transferred to tubes containing 750 µL of buffer Z (60 mM Na_2_HPO_4_; 40 mM NaH_2_PO_4_; 10 mM KCl; 1mM MgSO_4_) previously mixed with β-mercaptoethanol (ratio 100 mL buffer Z to 700 µL β-mercaptoethanol). The reaction tubes were then incubated at 37°C for 5 minutes. Following this, 250 µL of fresh ONPG (o-nitrophenyl-β-D-galactopyranoside) at 4 mg/mL was added to each tube, followed by a brief vortex. The tubes were then incubated at 37°C for 1 hour to allow the enzymatic reaction to proceed. After the incubation period, once the solutions turned yellow, the reaction was halted by adding 500 µL of 1 M sodium carbonate (Na_2_CO_3_) to each tube, followed by another 3-second vortex. The tubes were then centrifuged for 5 minutes at 13.500 g to pellet cell debris. Following centrifugation, the OD_414_ of the clear supernatant was measured to assess β-galactosidase activity. This repression assay was performed in biological triplicate for each condition.

### Growth curve measurements

Strains were revived from -80°C stocks by streaking onto LB plates containing Cm and Glc and incubated at 42°C overnight. The following day, colonies were inoculated in LB containing Cm and Glc at an OD_600_ of 0.1 and incubated at 42°C with shaking until an OD_600_ ∼0.6 was reached. Then, the cells were washed, diluted to obtain an OD_600_ of 0.003 in LB supplemented with 0.02% of Ara. Bacterial preparations were distributed in triplicates in 96-well microtiter plates and incubated at 30°C under agitation for 20h. Growth curves were monitored using a TECAN Infinite microplate reader with OD_600_ taken at 10-mins intervals. The growth curve assay was performed in biological triplicate for each strain.

### Plasmid stability assay

Strains were revived from -80°C stocks by streaking onto LB plates containing Cm and Glc and incubated at 42°C overnight. The following day, colonies were inoculated in LB containing Cm and Glc at an OD_600_ of 0.1 and incubated at 42°C with shaking until an OD_600_ ∼0.6 was reached. Then, the cells were washed, diluted to obtain an OD_600_ of 0.003 in LB containing Ara 0.02% (with or without the addition of Cm) and incubated with shaking at 30°C for approximately 10 generations, as estimated by optical density. To achieve growth for approximately 15 generations, cultures were back-diluted after 9 generations the same day and for 20 generations, cultures were back-diluted after 10 generations and incubated overnight. When the expected number of generations was achieved, cultures were diluted and plated onto LB plates supplemented with Glc (total population) or LB plates supplemented with Glc and Cm (plasmid-carrying population). Plasmid stability was calculated as the ratio of antibiotic-resistant clones. The plasmid competition assay was performed in biological triplicate.

### Conjugation assay

Overnight cultures were back-diluted in LB supplemented with 300 µM DAP and Glc and incubated at 42°C for the donor, or in LB and incubated at 37°C for the recipient cells until an OD_600_ ∼0.6-0.8 was reached. A donor-to-recipient ratio of 1/1 was used, and conjugation was performed using the filter mating procedure (28), at 42°C for 3h. Subsequently, samples were diluted and plated onto LB plates supplemented with Glc (total recipient population) or LB plates supplemented with Glc and Cm (transconjugants population). Conjugation frequencies were calculated as the ratio of transconjugants.

### Library design and construction

The library was designed by randomly choosing targets within the *V. cholerae* N16961 (NC_002505.1 for chromosome 1 and NC_002506.1 for chromosome 2) containing a proper NGG PAM. 11, 025 guides targeting *V. cholerae* chromosomes were designed, supplemented by additional 100 control random guides (Supplementary Table 1). The resulting library of 11, 125 gRNAs was synthesized by Twist Bioscience. The library was constructed as described in Rousset *et al*. (11). Pooled oligo extension was performed using KAPA HiFi DNA polymerase and the FR222 primer. The library was subsequently amplified through polymerase chain reaction (PCR) using KAPA HiFi polymerase (95°C for 3 min, 6 cycles of 98°C for 20 s, 60°C for 15 s, and 72°C for 20 s, with a final extension at 72°C for 10 min) with primers FR221 and FR222 and purified by gel extraction (GeneJET PCR purification, Thermo Scientific). Next, pT312 (*ccdB*^+^) was digested with BsaI-HF-v2 (New England Biolabs) and the vector was gel purified. The plasmid library was then assembled using the Gibson methodology (29).

For the library transformation in *E. coli*, T045 cells were grown in LB at 37°C to an OD_600_ ∼ 1. Cells were washed 4 times in ice-cold water and resuspended in ice-cold water. A total of 40 electroporations were carried out, with each electroporation using 0.5 µL of the dialyzed Gibson assembly product. Following a 1 h of incubation at 42°C, cells were plated on 80 large LB-Cm-Glc plates and incubated overnight at 42°C resulting in ∼ 3.10^6^ colonies. The following day, each plate was washed with LB supplemented with Cm and Glc and pooled. Plasmid library was extracted by maxiprep and transformed into β2163 *E. coli* conjugative strain. After electroporation, cells were incubated at 42°C and plated on 80 large LB-Cm-DAP-Glc-Km plates and incubated overnight at 42°C yielding ∼ 2.10^6^ colonies. The next day, each plate was washed with LB supplemented with Cm, DAP, Glc and Km and pooled.

For the transfer of the library in *V. cholerae* N16961, *V. cholerae* N16961Δ*ddmABC* was constructed using natural transformation. The library was then transferred by conjugation. Transconjugants were plated on 40 large LB-Cm-Glc plates and incubated overnight at 42°C resulting in ∼ 2.10^6^ colonies. The next day, each plate was washed with LB supplemented with Cm and Glc and pooled.

### Screen design

*V. cholerae* cells conjugated with the library were inoculated at an OD_600_ of 0.1 in LB-Cm-Glc and incubated at 42°C with shaking until an OD_600_ ∼0.6 was reached. Then, the cells were washed, diluted to obtain an OD_600_ of 0.003 in LB containing Ara 0.02% and incubated with shaking at 30°C. Simultaneously, the remaining culture at 42°C was used for plasmid extraction (Midiprep) to obtain the plasmid DNA at 0 generations. Upon reaching the 9^th^ generation, cultures were back-diluted to an OD_600_ of 0.024 and incubated at 30°C until 15 generations were reached. Plasmids were subsequently extracted using a midiprep kit, yielding the DNA library after 15 generations.

### Illumina sample preparation and sequencing

Library sequencing was performed as described by Cui *et al*. (10). The sequencing library was generated using two Kappa PCR reactions (Roche) with primers detailed in table 4. Starting with 100 ng of library plasmid, the first PCR (98°C for 5min; two cycles of 98°C for 30 s, 60°C for 1 min 30 and 72°C for 30 s; 72°C for 5min) was conducted in a 20-µL reaction with 4 pmol of each primer, adding the first index and the UMI (Unique Molecule Identifier). An exonuclease I (Thermo Scientific) treatment was performed, followed by a PCR Purification (AMPure XP, Beckman Coulter). The second PCR (98°C for 5 min; twelve cycles of 98°C for 30s, 60°C for 1 min 30 and 72°C for 30 s; 72°C for 5 min) was performed in a 30-µL reaction with 30 pmol of each primer, adding the second index and flow cell attachment sequences. Samples were pooled (41.3 fmol of each sample) and the resulting 354-bp PCR DNA fragments were gel extracted. The final library concentration was assessed using the DNA 1000 ScreenTape on a TapeStation (Agilent), diluted to 750 pM and sequenced using a NextSeq 2000 sequencing system (Illumina) using a custom protocol detailed by Cui *et al*., (10). On average, each sample yielded 4.5 million reads per sample, representing a coverage of ∼400X.

### Data analysis

Indexes were used to de-multiplex the data using a custom Python script. UMI were counted to remove potential PCR duplicates, resulting in the use of 92% to 96% of the reads depending on the sample. The reproducibility among experimental duplicates in our biological triplicates was very high, demonstrated by a median Pearson’s correlation coefficient r of 0.991. sgRNAs with fewer than 20 reads in the initial library (0 generations) were discarded in all the replicates (0.7% of sgRNAs). Sample normalization was performed by adjusting for sample size variation. This normalization was achieved by utilizing the read counts of 96 out of 100 control guides, as one control guide (control 83) was not present in the pool, and three other control guides exhibited an off-target effect (control 9, 81, 98) and were excluded from the analysis. Genes containing only 1 sgRNA were discarded from the analysis. For each gene, the median value of at least 2 or 3 sgRNA was calculated and used as the representative value for that gene. Paired-analysis was conducted to assess the comparison between the sample after 15 generations and at 0 generations.

For each gene, the log2FoldChange (log2FC) value was calculated using the formula : _log2FCgene *x* = log2(_mean reads replicate 1, 2, 3 15 generations gene 𝑥𝑥 +1_) where mean reads replicate_ mean reads replicate 1, 2, 3 0 generations gene 𝑥𝑥+1 1, 2, 3 15 generations gene *x* represents the average reads of gene *x* across three replicates at 15 generations, and mean reads 1, 2, 3 0 generations gene *x* represents the average reads of gene *x* across three replicates at 0 generations. An independent t-test was conducted to compare the fold change values between the two conditions, resulting in corresponding p-values.

## RESULTS

### Design of a dCas9 expressing vector optimized for *Vibrio cholerae*

Our objective was to develop a platform with an inducible dCas9 and a sgRNA library to target the entire set of 3, 715 annotated genes in the pandemic strain *V. cholerae* N16961. Previously, CRISPRi has been employed in *V. cholerae* C6706, utilizing a dCas9 integrated into the chromosome under the regulation of an anhydrotetracycline promoter (30). However, a leakage of the promoter in the absence of inducer was observed, leading to a 3 to 10-fold decreased in colony-forming unit (C.F.U) when using a sgRNA targeting an essential gene. In the context of constructing a sgRNA library, it is important to significantly reduce this leakage as it may lead to the depletion of sgRNAs targeting essential genes, thereby introducing a significant bias into the library.

Considering the effectiveness of CRISPRi in *E. coli*, our initial approach was to test the pFR56 plasmid, where the dCas9 is present on a p15A origin of replication and controlled by a 2, 4-diacetylphloroglucinol (DAPG)-inducible PhlF promoter (11). The random sgRNA in the vector was replaced by a sgRNA targeting the essential *rpsL* gene (*vc0359*) of *V. cholerae*. The two plasmids containing the random or the *rpsL* sgRNA were transformed into our *E. coli* donor strain and then transferred to *V. cholerae* through conjugation. After conjugation, cells were streaked on a selective medium. Despite the absence of *dCas9* inducer, a notable difference in cell viability was observed between the cells carrying a random sgRNA and those carrying the *rpsL* one (Fig. 1A). Additionally, a drop-test assay of *V. cholerae* containing the vectors was performed on medium with or without inducer. A decrease in cell viability of at least one log was observed for the cells carrying the *rpsL* sgRNA without inducer (Fig. 1B), indicating that the phlF promoter is leaky in *V. cholerae*, thus preventing the use of this promoter to construct the sgRNA library.

**Figure 1.**
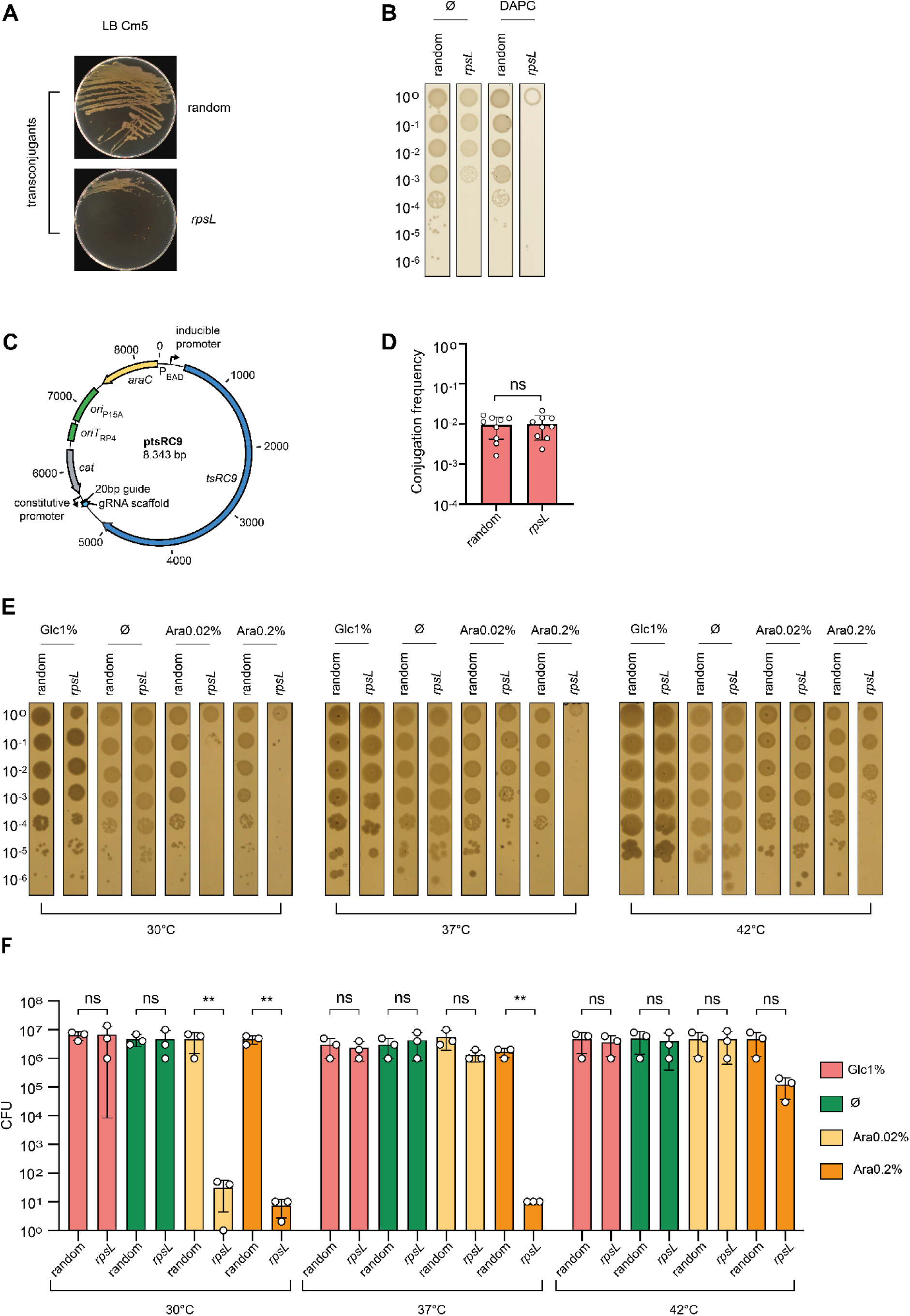
Development of an optimized CRISPRi vector for *Vibrio cholerae*. **A.** Conjugation plate assay depicting the result of conjugation between the donor *E. coli* strain ß-pFR56 carrying either a random guide or a guide targeting *rpsL* and *V. cholerae* strain on selective transconjugant medium (LB Cm5). **B.** Serial dilution drop test of *V. cholerae* harboring the plasmid pFR56 on LB agar with or without DAPG (50 μM). **C.** Plasmid map of ptsRC9 (pTN253), featuring the thermosensitive tsRC9 gene under the control of the P_BAD_ promoter (highlighted in yellow). The sgRNA (20bp guide + scaffold RNA) depicted in light blue is driven by a constitutive promoter. The chloramphenicol resistance gene (*cat*) is shown in gray. The plasmid includes a p15A origin of replication and an RP4 origin of transfer (both in green). **D.** Conjugation frequency of the CRISPRi vector with sgRNAs targeting essential or non-essential genes. Bar charts show the mean of conjugation frequency derived from nine independent experiments (n=9). Error bars show the standard deviation of mean (SD). Statistical comparisons (Student’s *t*-test, two-tailed) are as follow: ns, not significant; \*\**Pvalue < 0.01*. **E.** Serial dilution drop test of *V. cholerae* carrying the pTN253 plasmid with a random or *rpsL* sgRNA on LB medium (Ø) or supplemented with glucose 1% (Glc1%), or arabinose 0.02% - 0.2% (Ara). **F.** Bar charts showing the mean of the colony-forming unit (CFU) counts derived from three independent drop test experiments (n=3, individual plots) of *V. cholerae* with pTN253 carrying either a random or a *rpsL* sgRNA. Error bars show the standard deviation of mean (SD). Statistical comparisons (Student’s *t*-test, two-tailed) are as follow: ns, not significant; \*\**Pvalue < 0.01*. The different medium conditions are depicted: LB medium (green) or supplemented with glucose 1% (red), arabinose 0.02% (yellow), or arabinose 0.2% (orange).

To enhance the vector for *V. cholerae*, we replaced the dCas9 protein with tsRC9, a temperature-sensitive variant exhibiting robust activity at 30°C, but minimal activity at 42°C (19). Additionally, we replaced the PhlF promoter with the tightly regulated P_BAD_ promoter (Fig. 1C), known to be inducible with arabinose (Ara), and repressed with glucose (Glc) (24).

Comparison of the conjugation frequency, from *E. coli* to *V. cholerae*, between a plasmid harboring either a random sgRNA or a *rpsL* sgRNA revealed no significant difference, validating the tight control of the P_BAD_ promoter (Fig. 1D). This result confirmed that both sgRNAs targeting non-essential and essential genes can be transferred without introducing any differences at 42°C in the presence of glucose.

Serial dilution drop-test conducted with the new vector conjugated in *V. cholerae*, showed that cells harboring the *rpsL* sgRNA, regardless of the presence or absence of Glc, exhibited no difference in cell viability at 30 or 42°C, confirming minimal leakage from the P_BAD_ promoter (Fig. 1E, F). At 30°C, induction of tsRC9 with either 0.02% or 0.2% Ara led to growth arrest. However, at 37°C, growth arrest was only observed with 0.2% of Ara induction.

### Determination of effective gene silencing conditions

As a first step, we optimized the induction and temperature parameters of the system using a guide targeting *rpsL* (*vc0359*). However, it is important to note that *rpsL* is a housekeeping gene encoding the S12 ribosomal protein of the 30S subunit. Given its indispensable role in translation, we assumed that even a partial reduction in its expression could lead to a drastic reduction in cell viability. To ensure efficient gene silencing, we determined the strength of repression by our P_BAD_-tsRC9 system against the non-essential *lacZ* gene (*vc2338*), which is constitutively expressed in *V. cholerae* during exponential growth in rich medium (LB). We then performed a β-galactosidase activity assay to evaluate *lacZ* gene expression under different tsRC9 induction levels (0.02% or 0.2% Ara) over 15 generations of growth at 30°C. We observed that with a concentration of 0.02% or 0.2% Ara, the β-galactosidase activity was close to that of a Δ*lacZ* control strain. Specifically, the activity was reduced by approximatively 90% under both conditions (Fig. 2A). Thus, the level of repression achieved with 0.02% of Ara was considered high enough for our knockdown library screen.

**Figure 2.**
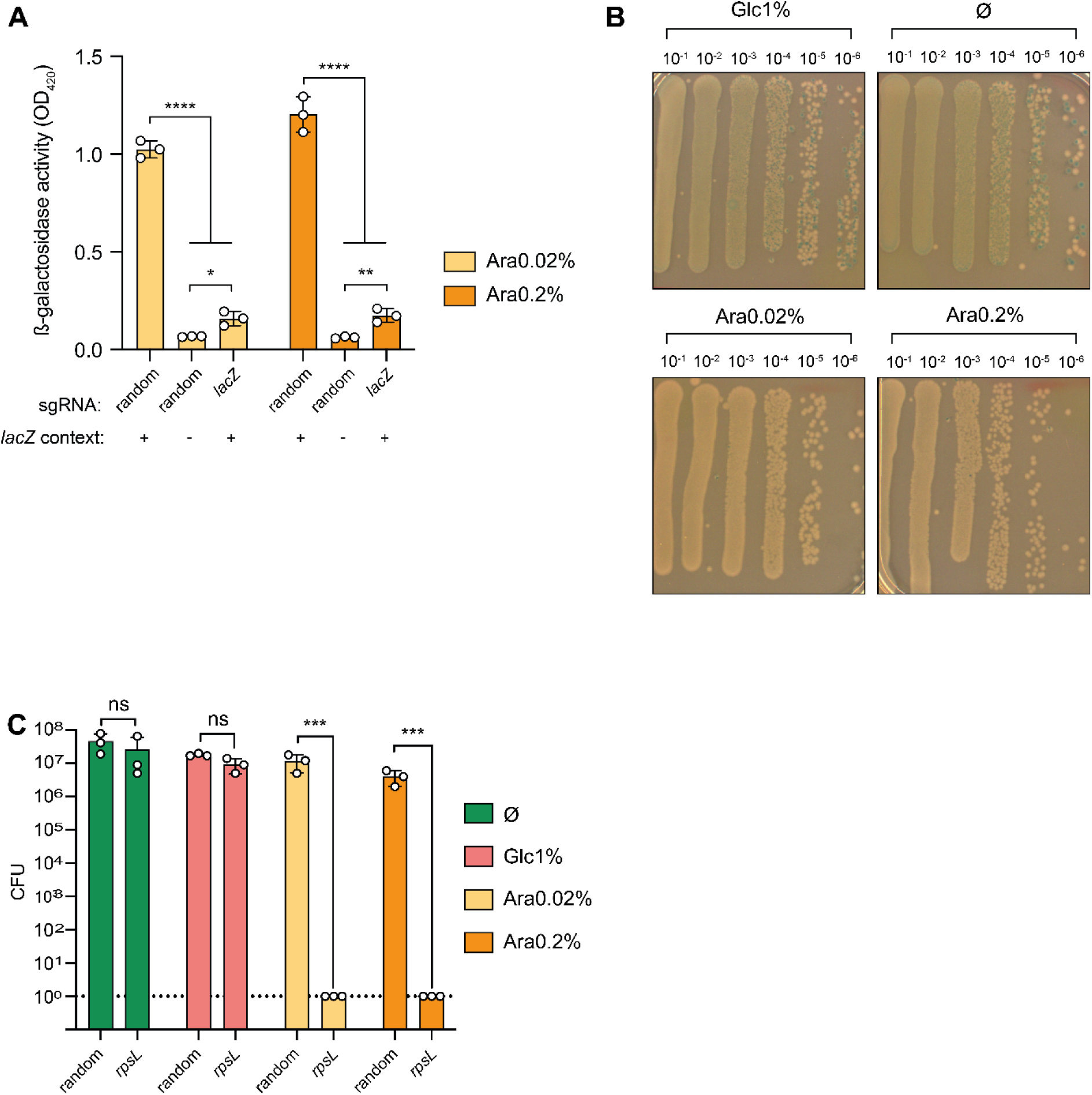
A plasmid containing a sgRNA targeting an essential gene is eliminated from the population. **A.** β-galactosidase assay reporting the expression of *lacZ* (*vc2338*) in *V. cholerae* strain carrying pTN253 with a random or a *lacZ* sgRNA. A Δ*lacZ* strain is also represented for basal ß-gal activity control. Two different tsRC9 induction conditions are depicted either with 0.02% or 0.2% arabinose. Bar charts represent the mean +/-s.d. from three independent experiments (n=3). Statistical comparisons (Student’s *t*-test, two-tailed) are as follow: ns, not significant; *, Pvalue < 0.05; **, Pvalue < 0.01; ****, Pvalue < 0.0001. **B.** Serial dilution drop test of a mixed population of *V. cholerae* carrying the pTN253 with a random (*lacZ*-, white colonies) or a *rpsL* sgRNA (*lacZ*+, blue colonies). The two strains competed during liquid culture for 10 generations in different tsRC9 induction conditions: LB glucose 1% (Glc1%), alone (Ø), arabinose 0.02% - 0.2% (Ara) as mentioned on top of the plates. **C.** Bar charts illustrating the colony-forming unit (CFU) counts derived from three independent competition assays (n=3) of *V. cholerae* with pTN253 carrying either a random or a *rpsL* sgRNA. Statistical comparisons (Student’s *t*-test, two-tailed) are as follow: ns, not significant; ***, Pvalue < 0.001.

Furthermore, we conducted a competition experiment to assess our system performance in a complex bacterial population. To this end, we initiated a competition between two bacterial strains: one, *lacZ*-, carrying a pTN253 with a random guide and the other, *lacZ+*, carrying a pTN253 with a guide targeting *rpsL*. These two strains were mixed and grown in liquid culture over 10 generations under different conditions (LB only, 1% Glc, 0.02% Ara, and 0.2% Ara). Induction of tsRC9 with either 0.02% or 0.2% Ara led to complete elimination of the population carrying the *rpsL* sgRNA (Fig. 2B, C), while in LB only or 1% Glc, there were no significant difference between the number of colony-forming unit (CFU), carrying either the *rpsL* or random sgRNA.

Altogether, these experiments demonstrate that our system fully meets our expectations of achieving high target gene repression.

### Assessment of tsRC9 toxicity and determination of efficient plasmid stability conditions

To avoid biases in our library due to a significant toxicity of dCas9, favoring the selection of inactivated mutants, we measured the intrinsic toxicity of tsRC9 in liquid culture. We performed a growth curve during induction with 0.02% Ara, with or without tsRC9, at 30°C, while a random sgRNA was constitutively expressed in both cases. Our results showed no significant effect of the presence of tsRC9 on the growth curve (Fig. 3A). Thus, tsRC9 does not exhibit toxicity in our system when induced with 0.02% Ara and in the presence of a constitutively expressed random sgRNA.

**Figure 3.**
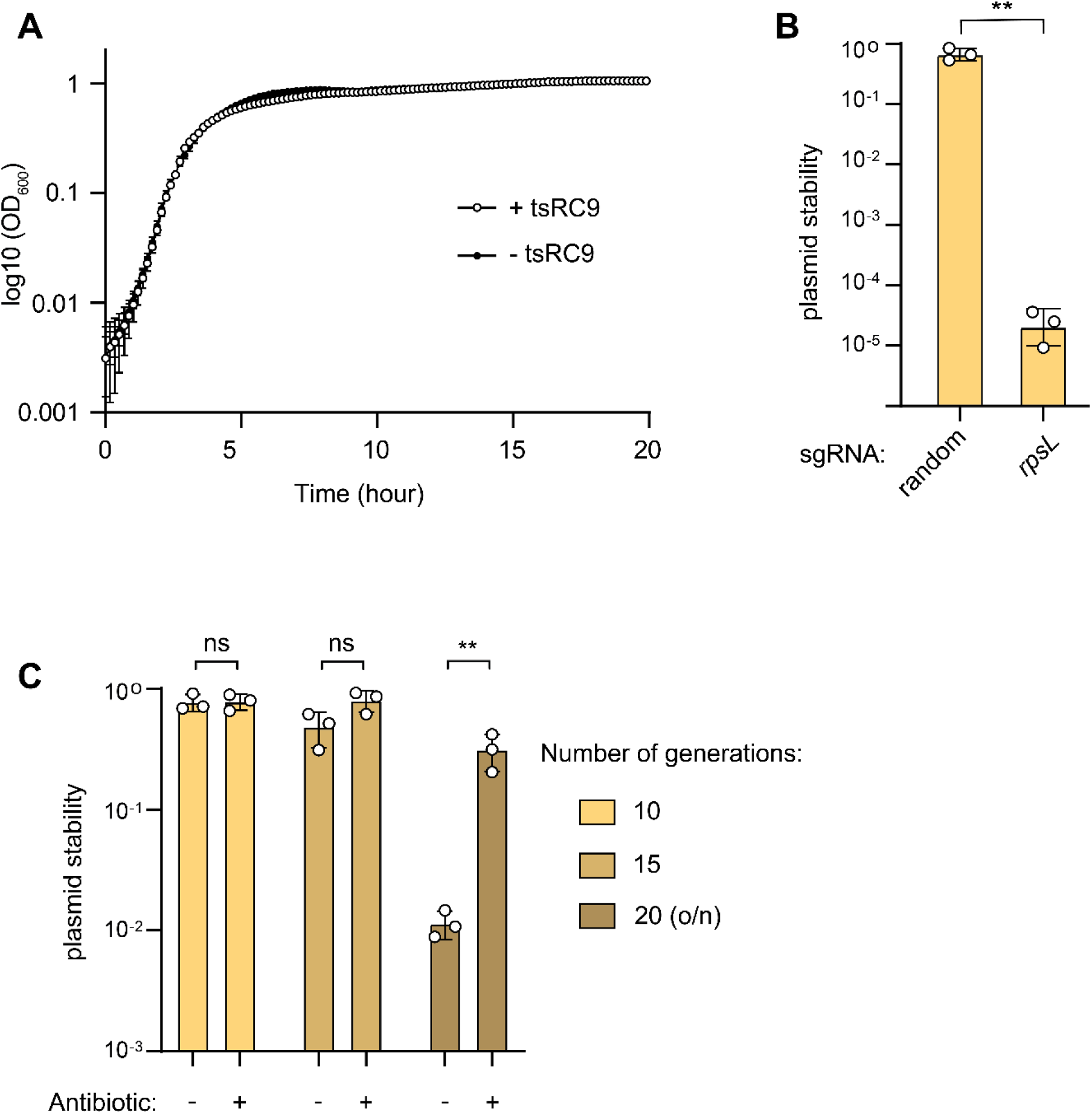
Stability of the CRISPRi vector in a Δ*ddmABC V. cholerae* strain. **A.** Growth curve in LB 0.02% arabinose of *V. cholerae* strain carrying a plasmid with (pTN253) or without the tsRC9 gene. **B.** Assessment of plasmid stability after 10 generations without antibiotic selection using different sgRNAs. Bar charts show the mean of plasmid stability derived from three independent experiments (n=3). Error bars show the standard deviation of mean (SD). Statistical comparisons (Student’s *t*-test, two-tailed) are as follow: \*\**Pvalue < 0.01*. **C.** Plasmid stability with or without antibiotic selection using a random sgRNA for 10, 15 or 20 generations. Bar charts represent the mean of plasmid stability derived from three independent experiments (n=3). Error bars show the standard deviation of mean (SD). Statistical comparisons (Student’s *t*-test, two-tailed) are as follow: ns: not significant, \*\**Pvalue < 0.01*.

A recent study revealed that *V. cholerae* El Tor strains harbor two defense modules, *ddmABC* and *ddmDE*, facilitating rapid plasmid degradation (22). Given that our CRISPRi vector carries a p15A origin of replication targeted by the *ddmABC* module but not by the *ddmDE* one, we removed *ddmABC* from our strain to prevent potential instability in the CRISPRi system. We tested the plasmid stability in this <u>Δ</u>*ddmABC* context. Our initial assessment involved testing the stability of a plasmid carrying either a random sgRNA or a *rpsL* sgRNA for 10 generations in 0.02% Ara. We observed that when the sgRNA did not target an essential gene, approximately 70% of the cells retained the plasmid (Fig. 3B). Conversely, targeting an essential gene resulted in the loss of the plasmid in almost all cells (Fig. 3B). This finding suggests that within our sgRNA library, those targeting essential genes lead to plasmid loss, resulting in a depletion of these sgRNAs in the pool.

Subsequently, we aimed at determining the maximum number of generations that could be performed without affecting plasmid stability. Using the plasmid carrying a random sgRNA, we assessed plasmid stability with or without antibiotic selection over a day for 10 and 15 generations, or with an overnight culture for 20 generations (Fig. 3C). We observed that for overday cultures, there was no significant difference in plasmid stability with or without antibiotic. However, with an overnight incubation to achieve 20 generations, there was a significant impact on plasmid stability, with only 1% of the cells retaining the plasmid. We first examined whether the *tsRC9* gene contributed to plasmid instability during prolonged stationary phases (Fig. S1A). Our findings demonstrated that while *tsRC9* did not affect plasmid stability during overday cultures, it was responsible for instability during overnight cultures (Fig. S1A). Next, we sought to understand the dynamics of tsRC9-containing plasmid instability (Fig. S1B). Our results demonstrate that, initially, cells begin to slowly lose the plasmid as they enter the stationary phase. However, extensive plasmid loss occurs over an extended period in the stationary phase, specifically after more than 7 hours through an unknown mechanism (Fig. S1B).

Consequently, our CRISPRi system cannot be utilized with overnight cultures but remains stable for up to 15 generations without antibiotic selection during an overday culture.

### Construction of the *Vibrio cholerae* sgRNA library and identification of essential genes

We designed a library targeting both chromosome 1 (NC_002505) and chromosome 2 (NC_002506) of *V. cholerae* N16961. A total of 11, 025 sgRNAs were designed to theoretically target 3, 706 genes out of 3, 715 (Supplementary Table 1). After cloning in *E. coli*, and conjugation to *V. cholerae*, the sequencing of the resulting library revealed a total of 10, 957 sgRNAs, enabling the potential silencing and analysis of 3, 674 genes.

To assess the functionality of our *V. cholerae* N16961 sgRNA library, we attempted to identify previously known essential genes in our screen by subjecting the library to 15 generations of growth in a rich medium (Fig. 4A). Throughout this experiment, sgRNAs detrimental to the bacterium’s fitness were expected to be depleted from the library. DNA from the sgRNA library was extracted both at 0 generations (before dCas9 induction) and at 15 generations (after dCas9 induction). Using the number of reads as measure of each guide’s abundance in the library, we calculated the log2 fold-change (log2FC) as a metric for relative gene fitness.

**Figure 4.**
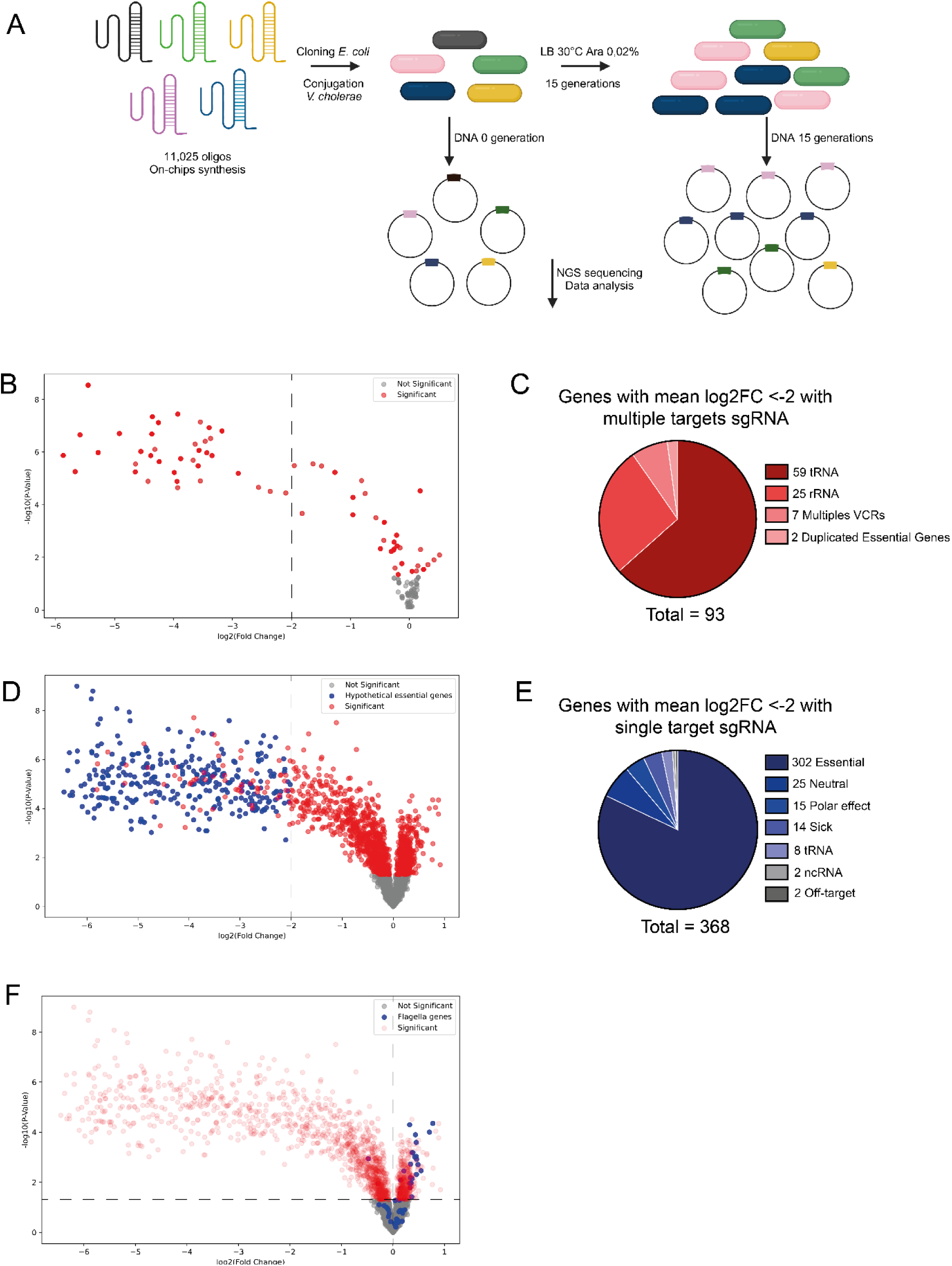
An arabinose-inducible CRISPRi system in *Vibrio cholerae*. **A.** Schematic overview of *V. cholerae* screen in rich media to identify essential genes. A sgRNA library was constructed on a plasmid expressing tsRC9 under the control of an arabinose-inducible promoter (pTN253) in *E. coli* and conjugated in *V. cholerae*. Plasmidic DNA was harvested before tsRC9 induction and after 15 generations of induction. Next-Generation Sequencing (NGS) was performed to obtain the distribution of each sgRNAs. **B.** Volcano plot visualization of the results from the multiple targets sgRNA screen. Genes that are not statistically significant are depicted in grey, while statistically significant genes are highlighted in red. Among the significant genes, candidate essential genes are those with a median log2FC < -2. **C.** 93 candidate essential genes (median log2FC < -2) identified from the multiple targets sgRNA screen. VCRs, *Vibrio cholerae* repeats. **D.** Volcano-plot visualization of the results from the single target sgRNA screen. Genes that are not statistically significant are depicted in grey, whereas statistically significant genes are highlighted in red. Among the significant genes, those considered as hypothetical essential genes are indicated in blue. **E.** 368 candidate essential genes identified from the single target sgRNA screen. Essential genes are those previously described as essential in other Tn-seq studies of *V. cholerae* (2, 3) or identified in a CRISPRi screening of a closely related bacterium, *V. natriegens* (18). “Sick” genes are those known to exhibit growth defects when mutated or deleted. Genes labeled as “Polar effect” are genes located upstream of essential or “sick” genes. “Off-target” genes are those with a strong off-target effect mediated by one of their sgRNAs. **F.** Highlighting the genes involved in flagellum production and regulation. Genes that are not statistically significant are depicted in grey, whereas statistically significant genes are highlighted in red. The genes involved in the flagellum biosynthesis are highlighted in blue.

In our library of 10, 957 sgRNAs, 5.8% (637/10, 957) target multiple loci on *V. cholerae* chromosomes, while 94.2% (10, 320/10, 957) target unique loci. The multiple loci targets correspond to known repeated sequences: tRNAs, rRNAs, duplicated integron gene cassettes, recombination sites (*Vibrio cholerae* Repeats, VCRs), transposases and duplicated genes.

Among these 637 multiple positions, we were able to analyze 221 genes (Supplementary Table 2, Supplementary Table 3). By choosing an arbitrary threshold of -2 log2FC to minimize false positive genes, 42.1% (93/221) exhibited a strong growth defect (log2FC < -2) (Fig. 4B). The analysis of these genes shown that the majority were involved in the protein biosynthesis pathway, including tRNAs, rRNAs, and the duplicated elongation factor *tufA* and *tufB* (Fig. 4C).

Among these 10, 320 unique positions, we were able to analyze 3, 453 genes (Supplementary Table 2, Supplementary Table 4). When silenced, 10.7 % of these genes (368/3, 453) exhibited a strong growth defect (log2FC < -2). The analysis of the 100 most depleted genes showed that these genes were associated to central cellular functioning including protein translation, DNA replication and cellular lipid metabolic process, all of which known as essential functions. We also observed that 25.1 % (867/3, 453) exhibited a mild growth defect (significant genes with log2FC between 0 and -2), and 64.2 % (2, 218/3, 453) exhibited no growth defect (non-significant genes or genes with log2FC > 0) (Fig. 4D). Notably, of the genes displaying an important growth defect, 82% (302/368) of them were identified as hypothetical essential genes in various Tn-seq studies of *V. cholerae* (2, 3) or in a CRISPRi analysis of a closely related bacterium, *V. natriegens* (18) (Fig. 4D). Out of the 66 remaining genes, 14 are known to hinder *V. cholerae* growth when inactivated by a transposon insertion (“sick phenotype” genes) (2) and 8 tRNAs as well as 2 non-coding RNAs (ncRNA) were identified. Furthermore, 15 of these genes exhibited a polar effect, resulting in the silencing of a downstream essential or “sick” gene (Fig. 4E). For example, in *V. cholerae*, the peptide deformylase *def* is not essential due to the presence of two non-essential paralogs, *vc0046* and *vca0150* (2). However, in our screen, silencing *vca0150* displayed no growth defect, while silencing *vc0046* induced a strong growth defect which can be explained by the downstream silencing of the essential single copy of methionyl-tRNA formyltransferase *fmt* gene (*vc0045*). Additionally, 2 genes (*vc2264* and *vc2655*) were affected by an important off-target effect of one sgRNA, resulting in a log2FC just above the threshold.

Finally, 25 genes, which were found to lead to a strong growth defect when silenced, were identified as neutral in a Tn-seq study (2). Of these 25 genes, 8 were also recognized as “growth-supporting” in a recent CRISPRi screening of *V. natriegens* (18). Additionally, 3 genes are part of toxin/antitoxin (TA) systems: *brnA* and *higA2* (antitoxins) and *brnT* (toxin). Surprisingly, these two antitoxins were classified as neutral in a Tn-seq study of *V. cholerae* C6706 (2). Since, it has been proven that their corresponding TA systems are active in *V. cholerae* N16961 (31), this explains why we identify them as essential genes. In the case of the *brnT* toxin, silencing of this gene also inhibits downstream expression of the *brnA* antitoxin, and we believe that this deregulation may result in toxicity. For the 14 remaining genes exhibiting a strong growth-defect, it is plausible that these genes could be classified as essential or “sick” genes only under our experimental conditions.

We also investigated genes known to confer a fitness advantage when inactivated. In the case of *V. cholerae*, it is well established that the flagellum is highly energy-demanding for the cell, and its inactivation is rapidly selected in evolutionary experiments (32). By examining the various genes involved in flagellum production and regulation (*vc2066-2069*, *vc2120-vc2144*, *vc2187-vc2208*), we detected that 52% (26/50) of them confer a slight fitness advantage (significant genes with log2FC >0) (Fig. 4F).

Finally, we analyzed the 20 TA systems present in our *V. cholerae* strain, with one located on chromosome 1 and 19 embedded within the sedentary chromosomal integron (SCI) on chromosome 2 (Fig. S2A) (31, 33, 34). Using our sgRNA library, we were able to investigate 18 out of the 20 TA systems, as one antitoxin (from TA13) was not targeted by our sgRNA library, along with the two antisense non-coding RNAs present in the *vca0495* toxin gene (from TA18). This library also enabled the study of six duplicated TAs present in the SCI (Fig. S2A). We found that silencing 8 out of 18 antitoxins induced a strong growth defect, while silencing 9 out of 18 antitoxins induced a mild growth defect (Fig. S2B, Supplementary Table 5). The varying impact on growth may be due to the differential effect of the toxins. For example, it has been shown that expressing *parE1*/*parE3* in *E. coli* resulted in less toxicity than expressing the *parE2* toxin, in agreement with our results (Fig. S2B, compare A7 to A2-A8) (31). Silencing the antitoxin of TA10 (*vca0422*) had no negative impact on the growth, consistent with the presence of an inactivated toxin (*vc0423*) due to an illegitimate recombination event with another cassette (31).

Overall, the list of essential genes obtained during exponential growth largely overlaps with known essential genes of *V. cholerae*. These results underscore the potential of our sgRNA library to investigate the majority of *V. cholerae* genes and highlight the effectiveness of CRISPRi screens in the pandemic strain of *V. cholerae*.

## DISCUSSION

Here we present a new CRISPRi set-up optimized for *Vibrio cholerae N16961 El Tor O1*, enabling the generation of global knockdown libraries. This new tool is ideal for large-scale genetic screens due to its straightforward setup, especially when compared to Tn-seq, which requires numerous processing steps.

Furthermore, the CRISPRi system does not require extensive sequencing depth, as only 1 million reads per sample are sufficient to yield high-quality results, providing approximately 100x coverage for each guide. In contrast, quality Tn-seq analysis of the *Vibrio cholerae* genome typically demands 5 to 10 million reads per sample. Therefore, the CRISPRi approach allows for highly multiplexed samples, which is considerable for studies involving numerous conditions and biological replicates.

Another advantage of CRISPRi over Tn-seq is its ability to investigate the role of essential genes in specific mechanisms which are typically absent in Tn-seq data. However, with CRISPRi, we can compare the essentiality of a housekeeping gene between two conditions, by comparing the loss kinetics of the associated guide at different time points. Thus, this approach may provide valuable insights into the dynamic role of these genes under varying experimental conditions.

In this study, we demonstrated the utility of our system for conducting genetic screens using 0.02% Ara induction at 30°C in LB medium. Importantly, this system also offers adaptability across a broad spectrum of induction conditions, such as being used at 37°C with 0.2% of Ara induction, or allowing for partial repression, a feature that can be advantageous in various experimental setups. Moreover, the system exhibits thermosensitivity, enabling rapid deactivation of dCas9 by simply raising the temperature to 42°C. This feature facilitates transient gene repression within a short timeframe, followed by swift restoration of normal gene expression levels. To validate the effectiveness of our pooled-library approach, our main objective was to identify known essential genes within the *V. cholerae* genome. Using our sgRNA library, we successfully identified 82% of the genes previously annotated as essential in separate Tn-seq and CRISPRi studies (2, 3, 18). However, one limitation of this system is its incompatibility with overnight cultures. Genetic screening must be carried out over a single day of experimentation, encompassing approximately 10 hours of culture for a *Vibrio cholerae* wild type strain to complete approximately 15 generations at 30°C. During prolonged stationary phases, the plasmid tends to be lost by a yet unidentified mechanism.

Now that we have successfully addressed the expression and toxicity issues associated with dCas9 in *V. cholerae*, we have developed a new user-friendly tool available for the research community. This tool will be of great interest for studying *V. cholerae* functioning and for identifying the gene landscape involved in response to various stimuli such as drug exposure or phage predation, and in gene transfer processes like conjugation and natural transformation, which contribute to the rapid adaptation of these bacteria.

## DATA AVAILABILITY

The data reported in this study are available in the European Nucleotide Archive (ENA). https://www.ebi.ac.uk/ena/browser/home; accession number ERR13166705. All the other data used are included within the published article and its additional files.

## SUPPLEMENTARY DATA

**Additional file 1: Table S1.** The list of gRNAs used in this study. The file is divided into three sheets as follows: the full list of gRNAs, the list of gRNAs targeting monocopy genes in the chromosomes of *V. cholerae* and the list of gRNAs and the list of gRNAs targeting multicopy genes in the chromosomes of *V. cholerae*.

**Additional file 2: Table S2.** The number of UMI counts detected for each sgRNA in each replicate, along with normalized data (sgRNAs with fewer than 20 reads in the initial library at 0 generations were discarded in all the replicates, and normalization between each replicate was performed by adjusting for sample size variation). The file is divided into four sheets as follows: UMI counts for the multiple targets analysis, UMI counts for the single targets analysis, normalized data for the multiple targets analysis, and normalized data for the single targets analysis.

**Additional file 3: Table S3.** Results of the CRISPRi screening for the multicopy genes of *V. cholerae*. The file is divided into six sheets as follows: the results of the CRISPRi screening for all the multicopy genes, the list of significant multicopy genes with a log2FC < -2 (hypothetical essential genes), the list of tRNAs among the hypothetical essential genes, the list of rRNAs among the hypothetical essential genes, the list of VCRs among hypothetical essential genes and the list of duplicated essential genes among the hypothetical essential genes. **Additional file 4: Table S4.** Results of the CRISPRi screening for the monocopy genes of *V. cholerae*. The file is divided into eight sheets as follows: the results of the CRISPRi screening for all the monocopy genes, the list of significant monocopy genes with a log2FC < -2 (hypothetical essential genes), the list of known essential genes among the hypothetical essential genes, the list of known neutral genes among the hypothetical essential genes, the list of genes that can be attributed to a polar effect among hypothetical essential genes, the list of known “sick” genes among the hypothetical essential genes, the list of tRNAs and non-coding RNAs among the hypothetical essential genes and the list of genes that can be attributed to an off-target of one of their sgRNA among the hypothetical essential genes.

**Additional file 5: Table S5.** Results of the CRISPRi screening for the 18 TAs analyzed in *V. cholerae*.

**Additional file 6: Figure S1.**

**Additional file 7: Figure S2.**

## AUTHOR CONTRIBUTIONS

K.D, T.N, D.B, Z.B and C.L designed the study. C.L and D.M obtained the funding. K.D, T.N, S.P, B.D performed the experiments. K.D, T.N and A.M analyzed the data and interpreted the results. K.D, T.N, C.L and D.M wrote the first version of the paper. All authors contributed to the editing of, and approved, the final manuscript.

## ACKNOWLEDGEMENTS

We would like to thank Andreas Möglich for the sharing of pFR097 plasmid (containing tsRC9). We would also like to thank Solange Miele, Depardieu Florence and François Rousset for their experimental advice and helpful discussions. Additionally, we thank Marie-Eve Kennedy-Val for the funding of Théophile Niault. Finally, we thank Busra Toktas for technical assistance.

## FUNDING

This work was supported by the Institut Pasteur, the Centre National de la Recherche Scientifique (CNRS-UMR 3525), the Fondation pour la Recherche Médicale (FRM Grant No. EQU202103012569), ANR Chromintevol (ANR-21-CE12-0002-01), ANR Jeunes Chercheurs [ANR-19CE12-0001], and by the French Government’s Investissement d’Avenir program Laboratoire d’Excellence ‘Integrative Biology of Emerging Infectious Diseases’ [ANR-10-LABX-62-IBEID].

## CONFLICT OF INTEREST

None declared.

---

**Figure S1.**
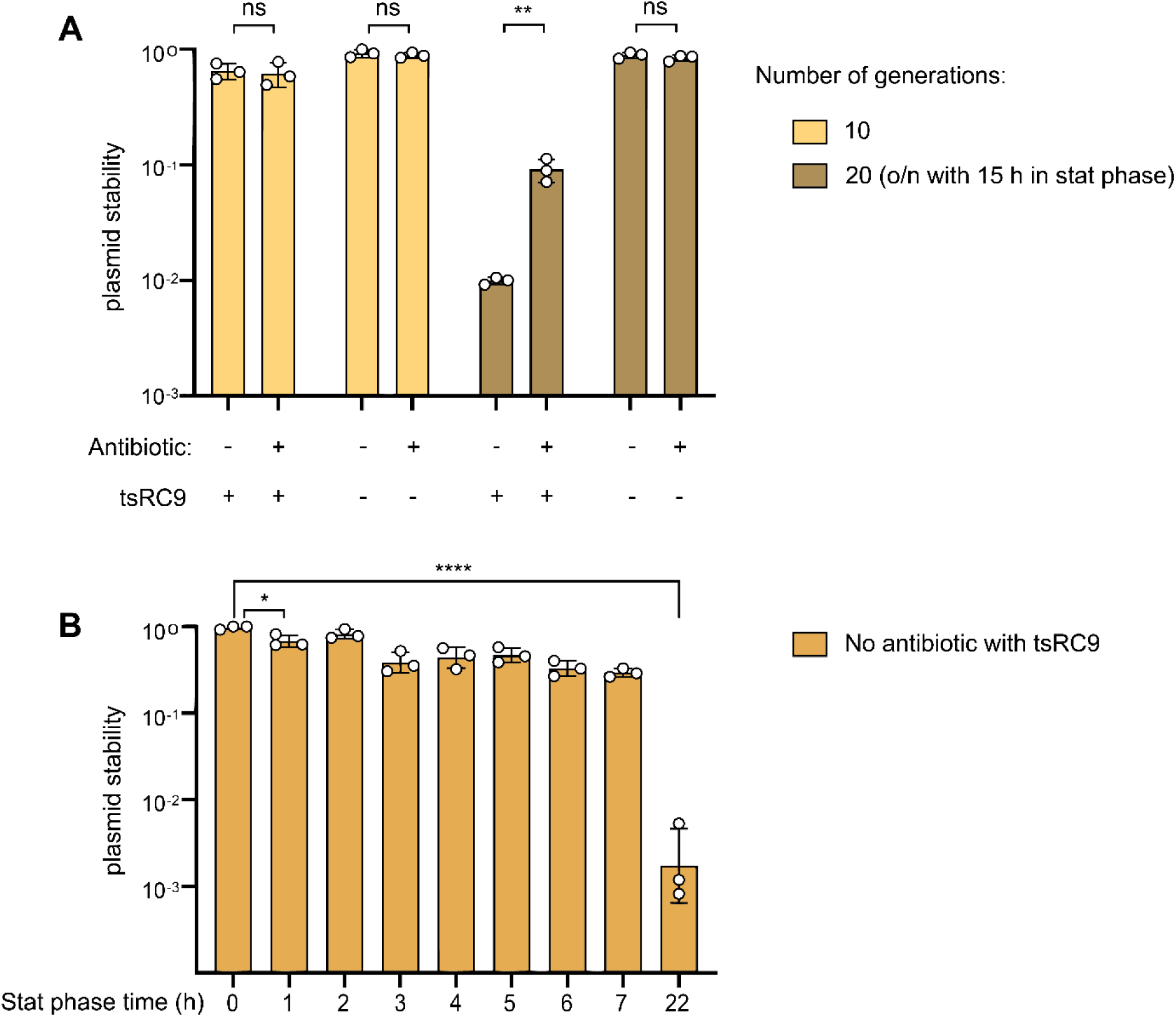
Identifying the factor causing plasmid instability in overnight cultures. **A**. Plasmid stability with or without antibiotic selection, with or without tsRC9 using a random sgRNA for 10 or 20 generations. Bar charts represent the mean of plasmid stability derived from three independent experiments (n=3). Error bars show the standard deviation of mean (SD). Statistical comparisons (Student’s *t*-test, two-tailed) are as follow: ns: not significant, \*\**Pvalue < 0.01*. **B**. Plasmid stability without antibiotic selection with tsRC9 using a random sgRNA during stationary phase. Bar charts represent the geometric mean of plasmid stability derived from three independent experiments (n=3). Error bars show the geometric standard deviation of mean (SD). Statistical comparisons (Student’s *t*-test, two-tailed) are as follow: ns: not significant, \**Pvalue < 0.05*, \*\*\*\**Pvalue < 0*.0001.

**Figure S2.**
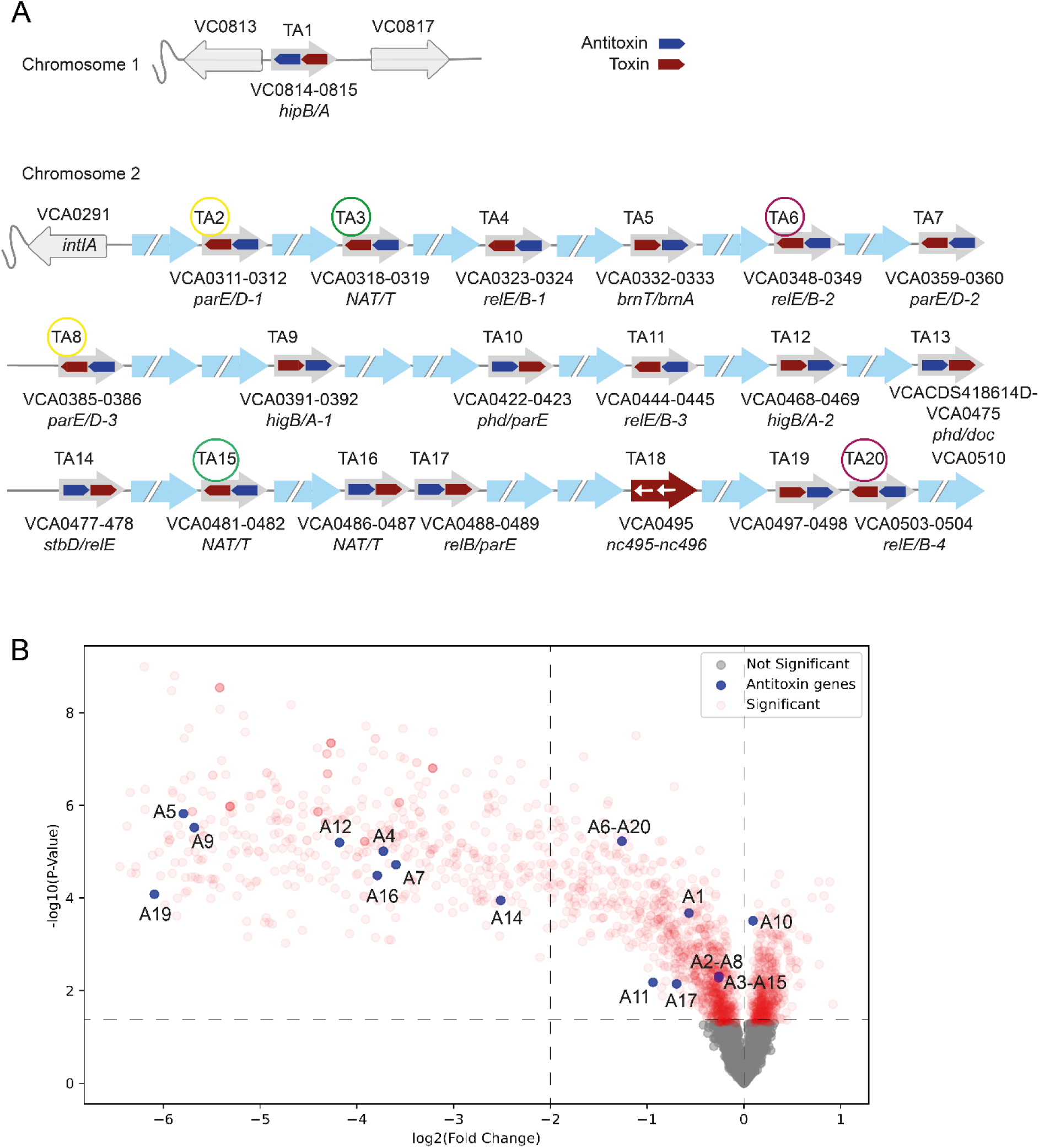
Validating CRISPRi screening through analysis of Toxin-Antitoxin systems in *V. cholerae*. **A.** Schematic representation of the 20 TAs in *V. cholerae.* Each TA is labelled with its name, a number, and orientation. White arrows indicate the presence of two non-coding RNAs within TA18. Duplicated TAs are depicted by circles of the same color. NAT/T stand for N-acetyltransferase/transcription factor. **B.** Volcano-plot visualization of the results from the single and multiple target sgRNA screen. Genes that are not statistically significant are depicted in grey, whereas statistically significant genes are highlighted in red. The antitoxins genes are highlighted in blue.

## Notes

### Competing Interest Statement

The authors have declared no competing interest.

